# Expanding the Orchidaceae Virome Through Analysis of Public-Domain Sequencing Data

**DOI:** 10.1101/2025.04.09.647952

**Authors:** Alejandra Pérez, Juliana Sánchez-Yali, Mónica Higuita, Adriana Ortiz-Reyes, Pablo A. Gutiérrez

**Affiliations:** Laboratorio de Microbiología Industrial, Facultad de Ciencias. Universidad Nacional de Colombia Sede Medellín. Carrera 65 Nro. 59A - 110, Medellín, Colombia; Departamento de Biociencias, Facultad de Ciencias. Universidad Nacional de Colombia Sede Medellín. Carrera 65 Nro. 59A - 110, Medellín, Colombia

**Keywords:** Orchidaceae, plant viruses, High-throughput sequencing, virus diversity

## Abstract

Orchidaceae is one of the largest plant families, renowned for its ornamental appeal, uses in food, cosmetics, and medicine, and its vital importance in ecosystem health and biodiversity. However, despite their diversity and economic significance, knowledge about the viruses associated with orchid species remains limited. Fortunately, the widespread adoption of high-throughput sequencing enables the collection of indirect information about the viruses associated with various plant hosts. In this work, we conducted an integrated data analysis of viruses associated with Orchidaceae by reexamining 2,789 public RNA-seq datasets encompassing 41 genera, 18 subtribes, and five subfamilies. We identified sequences associated with 163 viruses, 13 representing new associations with Orchidaceae. Additionally, 117 sequences likely represent putative novel species, including six tentative new genera within *Benyviridae*, *Betaflexiviridae, Phenuiviridae*, and *Tymoviridae*. Analysis of the updated Orchidaceae virome revealed a strong correlation between the pairs Alphaflexiviridae/Virgaviridae, Rhabdoviridae/Partitiviridae, and Partitiviridae/Secoviridae and a negative correlation between *Alphaflexiviridae*/*Partitiviridae*; *Virgaviridae/Partitiviridae*; *Rhabdoviridae*/*Alphaflexiviridae*; *Betaflexiviridae/Alphaflexiviridae*; and *Secoviridae*/*Alphaflexiviridae* in addition to correlations between virus genera and orchid lifestyles. A network analysis of the host range of orchid viruses was also performed in addition to a preliminary estimate of the size of the Orchidaceae virome.

**IMPORTANCE:** Despite their diversity, little is known about viruses associated with orchids. Plant viruses significantly threaten agriculture and may harm orchid conservation. Without detailed characterization of the orchid virome and suitable diagnostic tools, viruses could unintentionally enter ecosystems and cultivation fields from contaminated sources. Sequence information is crucial for developing detection tests for quick virus identification, supporting disease management strategies, and producing virus-free plants. In addition, understanding ecological factors contributing to virus transmission—like primary infection sources, vectors, and alternative hosts—is essential for selecting control measures to prevent viral diseases and manage outbreaks. Data mining provides a cost-effective method to bridge knowledge gaps, highlight trends that are useful for plant virus surveillance, and inform epidemiological decisions. The analysis presented here offers a global perspective on the Orchidaceae virome that could guide future research and virus vigilance programs and enhance our understanding of plant virus diversity.

## INTRODUCTION

With over 800 genera and more than 30,000 recognized species, Orchidaceae is the second largest plant family (Freudenstein, 2014; Pérez-Escobar et al., 2024; Wang et al., 2024). Orchids are primarily recognized as ornamentals, with species from the genera *Cattleya, Cymbidium, Dendrobium, Paphiopedilum*, and *Phalaenopsis* making up approximately 10% of the international fresh-cut flower market (Hinsley et al., 2018; Yuan et al., 2021; Zhang et al., 2022). Apart from their aesthetic appeal, orchids have a rich history of use in food, cosmetics, and traditional medicine (Bazzicalupo et al., 2023; Bulpitt, 2005; Castillo-Pérez et al., 2024; Fonge et al., 2019; Hadi et al., 2015). In the food industry, the most notable example is vanilla (*Vanilla*), which is one of the most expensive spices in the world and is expected to reach USD 4.3 billion in the global market by 2025 (da Silva Oliveira et al., 2022; Khan et al., 2022). Other orchids from the genera *Anacamptis, Dactylorhiza, Disa, Habenaria, Himantoglossum*, and *Satyrium* are popular ingredients in drinks and desserts within local cuisines throughout Africa, Asia, and some European countries (Bingham, 2009; Decary, 1955; Ghorbani et al., 2017; Hinsley et al., 2018; Kaputo, 1996; Kasparek & Grimm, 1999; Kreziou et al., 2016). In addition, dozens of orchid species, mostly geophytes, have been widely used in traditional medicine for centuries, especially in China, India, and parts of Africa (Bazzicalupo et al., 2023; Bulpitt et al., 2005, 2007; Dhyani et al., 2010; Hinsley et al., 2018; Kong et al., 2003; Randriamihariso et al., 2015), and there is growing interest within the cosmeceutical industry in the formulation of perfumes, colognes, body lotions, and soaps (Hsiao et al., 2011; Yang et al., 2022; Zhu et al., 2013). More importantly, Orchids are essential for ecosystem health and biodiversity (Vitt et al., 2023). They support pollinators, maintain soil health, and indicate environmental changes such as deforestation and pollution (Fay et al., 2018; Zhang et al., 2018). While they are not direct food sources, they feed pollinators that benefit fruit-bearing plants (Nilsson, 1992; Perkins et al., 2023). Additionally, orchids rely on mycorrhizal fungi for seed germination and growth, enhancing soil health and microbial diversity (Fabvre-Godal et al., 2020; Zhang et al., 2018).

Despite their diversity, orchids have been reported to be infected with only a few dozen viruses, the most important of which are cymbidium mosaic virus (CymMV, *Potexvirus cymbidii*) and odontoglossum ringspot virus (ORSV, *Tobamovirus odontoglossi*) (Chen et al., 2019; Koh et al., 2014; Ryu et al., 1995; Wong et al., 1997). These two viruses are highly prevalent worldwide and can infect a wide range of species across several genera, causing significant losses due to stunted growth and reductions in flower size and quality (Ajjikuttira et al., 2009; Chen et al., 2019; Koh et al., 2014; Soto-Valladares et al., 2012; Sun et al., 202; Yusop et al., 2022). Other common orchid viruses include orchid fleck virus (OFV, *Dichorhavirus orchidaceae*), and vanilla mosaic virus (VanMV; *Potyvirus dasheenis*) (Farreyrol et al., 2006; Peng et al., 2013; Puli’uvea et al., 2017; Randriambololona et al., 2022). OFV is a virus of a worldwide distribution that infects a wide range of orchid genera and ornamentals from other families (Mei et al., 2016; Otero-Colina et al., 2021; Read et al., 2021) and can also infect citrus species, causing the citrus leprosis (CL) disease (Cook et al., 2019; Dietzgen et al., 2016; Roy et al., 2014, 2015). VanMV, on the other hand, is a dasheen mosaic virus (DsMV) strain that infects vanilla and other edible and ornamental orchids (Babu, 2014; Jordan et al., 2002; Farreyrol et al., 2006; Puli’uvea et al., 2017). In addition, viruses from other plant families are often found to infect orchids, as evidenced by frequent reports of infection with bean common mosaic virus (BCMV, *Potyvirus phaseovulgaris*), cucumber mosaic virus (CMV, *Cucumovirus CMV*), watermelon mosaic virus (WMV, *Potyvirus citrulli*), turnip mosaic virus (TuMV, *Potyvirus rapae*), impatiens necrotic spot virus (INSV, *orthotospovirus impatiensnecromaculae*), tomato spotted wilt virus (TSWV, *Orthotospovirus tomatomaculae*), capsicum chlorosis virus (CaCV, *Orthotospovirus capsiciflavi*); tobacco rattle virus (TRV, *Tobravirus tabaci*), and tomato aspermy virus (TAV, *Cucumovirus TAV*), among others (Baker et al., 2007; Grisoni et al., 2004; Madhubala et al., 2005; Nguyen et al., 2013; Yoon et al., 2022; Zhang et al., 2010). If we assume that each plant species can host at least one virus, it is evident that our knowledge of orchid viruses is far from complete (Anthony et al., 2013; Carroll et al., 2018; Paez-Espino et al., 2016; Solé & Elena 2019). This hypothesis is supported by sequencing surveys conducted in Australia, Europe, and Asia, which have uncovered numerous novel viruses in a limited number of orchid samples (Chao et al.,2022; Wylie et al., 2012; Wylie et al., 2013a,b; Komínek et al., 2019; Huang et al., 2019; Kondo et al., 2014).

Viruses also endanger orchid conservation efforts. It has been estimated that around 30% of orchid species are at risk of extinction (Fay, 2018; Swarts & Dixon, 2009; Vitt et al., 2023), and one method for protecting these endangered species involves cryo collections and *in vitro* propagation (Das et al., 2021; Merrit et al., 2014; Popova et al., 2016). Unfortunately, without a detailed characterization of the orchid virome and the availability of suitable diagnostic tools, viruses may be unintentionally introduced into natural ecosystems from contaminated propagation sources (Campol et al., 2024; Das et al., 2021; Wylie et al., 2013). In this context, sequence information plays a crucial role by aiding in the development of detection tests for the prompt identification of viruses, thereby supporting integrated disease management strategies and the production of virus-free plants (Kalimuthu et al., 2022; Katsarou et al., 2019; Sun et al., 2023; Tatineti et al., 2023). Additionally, understanding the ecological factors that contribute to virus transmission—such as primary infection sources, involved vectors, and alternative hosts—is essential for choosing suitable control measures that prevent or mitigate the emergence of viral diseases and manage potential outbreaks (Elena et al., 2014; Jones et al., 2009; Lefeuvre et al., 2019). Traditionally, gathering epidemiological information on a global scale can pose significant logistical challenges (Buckee, 2020). Fortunately, the large volume of publicly available high-throughput sequencing data can serve as an indirect, cost-effective method for obtaining indirect epidemiological data and provide valuable genomic insights into new viruses, variants, and virus-host relationships involving plants of interest (Debat et al. 2023; Higuita et al. 2024ab; Sidharthan et al. 2022, 2024). Here, we present a meta-analysis of the Orchidaceae virome that integrates knowledge from published records, annotated sequence data, and data mining of public RNA-seq data that expands our current knowledge of orchid-associated viruses.

## METHODS

### RNAseq data

Reads were obtained from the Sequence Read Archive (SRA) at https://www.ncbi.nlm.nih.gov/sra and downloaded using the fasterq-dump tool from sratoolkit 3.0.5. We inspected 2,789 Orchidaceae datasets from 271 Bioprojects comprising 203 orchid species or hybrids (Supplementary Data). A comprehensive overview of sequencing technologies, source tissue, and library preparation methods can be found in the supplementary material and is illustrated in Supp. Fig.1. Datasets were assembled with rnaSPADES (Bushmanova et al. 2019). Putative viral sequences were identified using DIAMOND (Buchfink et al. 2015) with a database that included all reference viral proteins from NCBI (683,315 sequences), all known proteins from the family Orchidaceae (106,405 sequences), and the EnsembleRapid dataset (2,888,407 sequences), which features a non-redundant collection of proteins from all cellular organisms (Martin et al. 2023). The reported phylogenetic placement of hosts was confirmed through phylogenetic analysis of concatenated *matK* and *rbcL* sequences (CBOL 2009; Hollingsworth et al. 2011, 2016; Kress et al. 2005). Only viruses associated with families known to infect plants were included in this analysis; a complete list of additional suspected viral sequences is provided in the supplementary material. To reduce the likelihood of false associations, datasets that included *matK* or *rbcL* sequences from non-orchidaceous hosts were excluded from this work (Suppl. Data). Selected viral sequences were verified manually and inspected for the presence of the expected molecular features reported by the ICTV (Lefkowitz et al. 2018), following the procedures detailed by Higuita et al. (2024a,b). Arthropod and nematode associations were determined through analysis of *coxI* barcodes (Palomares et al., 2017; Salis et al., 2024; Wilson, 2012) and provided in the supplementary data. All sequences discussed in the text are provided as supplementary data and labeled with the corresponding Bioproject and SRA accession codes.

### Phylogenetics

Trees were calculated using the maximum-likelihood method using IQ-TREE (Nguyen et al. 2015). Support values were estimated using a likelihood ratio test with 10,000 replicates and the ultrafast bootstrap approximation UFBoot with 1,000 replicates (Minh et al., 2013). Multiple sequence alignments were generated with MAFFT (Katoh et al., 2002) and edited to remove incomplete ends and ambiguous positions using JalView (Waterhouse et al., 2009). Evolution models were selected using ModelFinder (Kalyaanamoorthy et al. 2017).

### Virus associations

The enrichment or depletion of viruses relative to the taxonomic rank of hosts or the presence of other viruses was analyzed using Fisheŕs exact test. The relative proportion of each virus was compared to the proportion observed for the complete virome. The transmission mechanism for each virus was obtained from the virus plant virus transmission database (Peters et al., 2024). Enrichment analyses were performed using Fisheŕs exact test implemented in the Python package SciPy (Virtanen et al., 2020). Networks were built with the Python package NetworkX (Hagberg et al. 2008). Network communities were identified using the Louvain algorithm (Blondel et al. 2008). Closeness and betweenness centrality were measured using the Wasserman and Faust formula (Wasserman and Faust, 1994) and the Ulrik Brandes algorithm (Brandes 2001, 2008; Freeman 1977). Networks were visualized and rendered with Gephi (Bastian et al., 2009) and are provided as supplementary data.

### Rarefaction analysis

For the rarefaction analysis, we generated a set comprising all the associations between viruses and orchid genera. Each point corresponded to the average number of viruses found after sampling between 1 and 100 virus associations, using ten repetitions in each trial. Results were fitted to negative exponential and sigmoidal models (Colwell and Coddington, 1994) using SciPy (Virtanen et al., 2020).

### Data availability

All data from this study is available at Zenodo 10.5281/zenodo.15113200

## RESULTS

### Global description of results

After inspecting 2,789 RNA-seq datasets, we identified 163 putative viruses associated with 114 orchid species or hybrids from 41 genera, 18 subtribes, and five subfamilies (Fig 1, Supplementary Fig. 1C and D). Excluding unidentified species and hybrids, we found an average of 1.6 viruses per orchid species (Supp. Fig.2A). Fisher’s exact test shows that this ratio applies to most genera, except *Phalaenopsis* (0.42 ratio, *P*=1.38e-5) and *Cymbidium* (0.86, *P*=0.007), which had fewer viruses than expected. In the subfamily Apostasiodeae, the only virus detected was CymMV, which was found in *Apostasia wallichi*, *Neuwiedia malipoensis*, and *N. zollingeri*. In Vanilloideae, we identified three viruses: vanilla latent virus (VLV, *Allexivirus latensvanillae*) in *V. planifolia*, CymMV in *Vanilla shenzhenica,* and a new *Deltapartitivirus* in *Galeola faberi*. In the subfamily Cypripedioideae, we identified 35 viruses in datasets from the tribes Cypripedieae (*Cypripedium*), Phragmipedieae (*Paphiopedilum*, *Phragmipedium*), and Mexipedieae (*Mexipedium*). In this subfamily, the highest number of viruses was observed in the genus *Paphiopedilum*, totaling 21 virus species across eight genera, followed by *Cypripedium*, with ten viruses from seven families (Fig. 1A and B). The tribe Cypripedioideae was also enriched with viruses from *Secoviridae* (*P*=8.4×10^-17^) and *Partitiviridae* (*P*=5.8×10^-5^) (Supplementary Fig 2B). Most of our findings occurred in the subfamilies Orchidoideae and Epidendroideae (Fig. 1A). Our analysis of Orchidoideae included 32 species from 15 genera in the tribes Cranichidae, Cymbidieae, Diurideae, and Orchideae. Orchidaeae was enriched in viruses from *Partitiviridae* (*P*=6.5×10^-17^), while the tribe Diuridae was also found to be enriched with members of the *Aspiviridae* (*P*=1.2 x10^-5^). The genera with the largest diversity of viruses were *Gymnadenia* (13 viruses), *Platanthera* (10 viruses), and *Dactylorhiza* (9 viruses). Lastly, in the subfamily Epidendroideae, our analysis included 57 orchid species from 17 genera in the tribes Arethuseae, Calypsoeae, Collabieae, Cymbidieae, Epidendreae, Gastrodieae, Malaxideae, Neottieae, and Vandeae (Fig 1A). This subfamily was enriched in *Virgaviridae* (*P*=3.8 x10^-5^) and *Alphaflexiviridae* (*P*=2.9 x10^-10^). At the tribe level, Neottieae was enriched with *Benyiviridae* (*P*=0.0002) and Podochileae with *Potyviridae* (*P*=3.3 x10^-10^). The genera with the most viruses were *Dendrobium* (21 viruses) and *Epipactis* (12).

**Figure 1.**
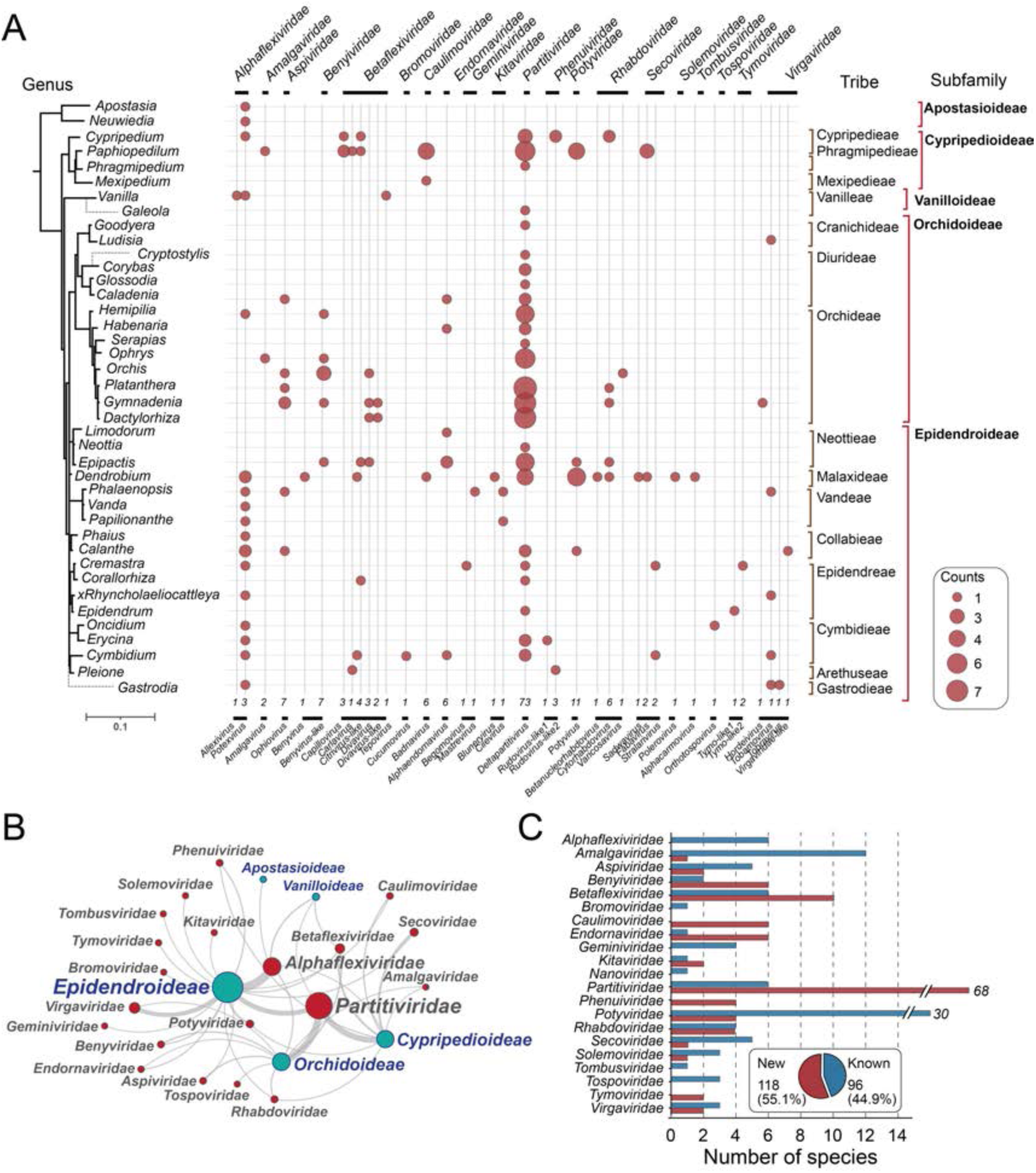
Orchid viruses found in public RNA-seq data. A) Correlation plot illustrating the number of virus families and genera detected in different orchid genera. The phylogenetic tree to the right illustrates the evolutionary relationship between the orchids analyzed in this study. The tree was based on *rbcL*/*matK* genes. The dotted lines correspond to genera for which *rbcL*/*matK* sequences were not available or were of insufficient length. B) Bipartite network illustrating the associations between virus families (blue) and orchid subfamilies (red). Edge thickness indicates the number of associations, while node size reflects the number of viruses found. C). Barplots illustrating the number of new virus species detected by RNA-seq and the known orchid viruses found in the NCBI virus database.

Regarding viral sequences, we identified 117 (74.2%) putative new viral species from fifteen virus families, including six tentative new genera within *Benyviridae*, *Betaflexiviridae*, *Phenuiviridae*, and *Tymoviridae* (Fig 1A and C; Supplementary Data). Among the known viruses, thirteen correspond to new host associations with Orchidaceae: barley yellow striate mosaic virus (BYSM, *Cytorhabdovirus hordei*), beet necrotic yellow vein virus (BNYVV, *Benyvirus necrobetae*), broad bean wilt virus 2 (BBWV2, *Fabavirus betaviciae*), cactus virus X (CVX, *Potexvirus ecscacti*), chickpea redleaf virus 2 (CpRLV2, *Mastrevirus rubrumsecundi*), east asian passiflora virus (EAPV, *Potyvirus orionspassiflorae*), grapevine fabavirus (GFabV, *Fabavirus vitis*), lily mottle virus (LMoV, *Potyvirus lilimaculae*), potato Virus H (PotVH, *Carlavirus chisolani*), potato virus Y (PVY, *Potyvirus yituberosi*), shallot yellow stripe virus (SYSV, *Potyvirus ascaloniae*), sugarcane mosaic virus (SCMV, *Potyvirus sacchari*), and tomato yellow leaf curl virus (TYLCV, *Begomovirus coheni*). A detailed description of findings within each virus family is provided in the following sections.

***Alphaflexiviridae.*** We identified four *Alphaflexiviridae* from the genera *Potexvirus* and *Allexivirus*: CymMV, CVX, phaius virus X (PhVX, *P. ecsphaii*), and vanilla latent virus (VLV, *A. latensvanillae*). CymMV is one of the most prevalent viruses in orchids (Yusop et al., 2022). This observation agrees with our results, as CymMV was found in 217 datasets from 36 species and hybrids from the genera *Apostasia, Cremastra, Cymbidium, Cypripedium, Dendrobium, Erycina, Gastrodia, Hemipilia, Neuwiedia, Oncidium, Papilionanthe, Phaius, Phalaenopsis, Vanda,* and *Vanilla* (Figure 1, Supp. Fig.3A). CymMV was highly prevalent in the subfamily Epidendroideae but rare in Orchidoideae (Fig 1A). In agreement with previous studies (Moles et al., 2007), phylogenetic analysis of CP sequences revealed two CymMV clades with nucleotide sequence identities of ∼97% (Clade I) and ∼87% (Clade II) to the reference CymMV sequence (NC_001812) (Supp. Fig.3A). PhVX was detected in *Dendrobium nobile,* with sequences sharing nucleotide identities in the 76.2-79.3% range to the reference sequence NC_010295 from the orchid *Phaius flavus* (Kawakami et al., 2008). VLV was detected in *V. planifolia* with nucleotide identities in the 92.9-94.5% range to the reference sequence NC_035204, also from *V. planifolia* (Grisoni et al., 2017). Finally, CVX was associated with *Calanthe tsoongiana* and was most closely related to an isolate from alfalfa (BK011045, 97.4%) (Jiang et al., 2019) (Supp. Fig.3A). To the best of our knowledge, this would represent the first report of CVX in orchids.

### Amalgaviridae

We identified ophrys fusca amalgavirus 1 (OfAV1) in *Ophrys iricolor* and a sequence of 3,042 nt in *P. armeniacum* that likely represents a putative new species (Supp. Fig 3B). This sequence was ∼66% identical at the amino acid level to epipogium amalgavirus 1 and bletilla striata amalgavirus 1. For this new virus, we propose the name paphiopedilum armeniacum-associated amalgavirus 1 (PaaAV1).

### Aspiviridae

We confirmed the presence of several orchid-infecting ophioviruses: caladenia ophiovirus (CalOV), gymnadenia ophiovirus den (GymOV-den), gymnadenia ophiovirus odo (GymOV-odo), dactylorhiza hatagirea ophiovirus (DacthOV), and phalaenopsis ophiovirus (PhalaOV) (Sidhartan et al., 2021; Debat et al., 2023) (Supp. Fig 3C). We also found two putative new *Ophiovirus* species in *Platanthera guangdongensis* and *Calanthe triplicata,* which we named platanthera-associated ophiovirus 1 (PgaOV1) and calanthe-associated ophiovirus 1 (CalanOV1). At the nucleotide level, PgaOV1 was most closely related to GymOV-den/odo (∼67.4-68.4%) and DacthOV (∼66.2%); CalanOV1 was most closely associated with PhalaOV (∼70.0%).

### Benyviridae

Dactylorhiza hatagirea beny-like virus (DhBLV) is the only Benyvirus-like virus reported in orchids (Sidharthan et al., 2021). In addition to DhBLV, we identified RNA2 and RNA3 segments of beet necrotic yellow virus (BNYVV*, Benyvirus necrobetae*) in *Dendrobidum officinale*, which would be the first *Benyvirus* in Orchidaceae (Figure 2A). We also found six new beny-like viruses related to DhBLV for which we propose the names ophrys fusca associated beny-like virus (OfaBLV1), hemipilia forrestii associated beny-like virus (HfaBLV1), orchis militaris associated beny-like virus (OmaBLV1), orchis hatagirea associated beny-like virus (OhaBLV1), gymnadenia rhellicani associated beny-like virus (GraBLV1), and epipactis helleborine associated beny-like virus (EhaBLV1) (Figure 2A). These beny-like viruses form a clade sister to the genus *Benyvirus,* supporting the suggestion that DhBLV is a member of a new taxon (Sidharthan et al., 2021). This claim is further supported by a cross-comparison of RdRp protein segments, which revealed identities in the 22.7-25.8% range with *Benyvirus* and 64.7%-78% between different species in the Beny-like clade (Figure 2B). These Beny-like sequences were ∼7.4-7.9 kb long with a poly-A tail. The genomic RNA encoded an RdRp protein of ∼260kDa with a viral RNA helicase (PF01443) domain in the N-terminus (884 – 1120) and an RNA-dependent RNA polymerase (PF00978) at the C-terminus (1978 – 2294). In contrast to members of the genus *Benyvirus*, the RdRp protein lacked methyltransferase and papain-like protease motifs (Gilmer et al., 2017). The RdRp has two non-cytoplasmic domains corresponding to the Helicase and RdRp domains and three transmembrane regions (Fig 2C). ORF2 encodes a putative polytopic membrane protein of ∼17 kDa, with a non-cytoplasmic domain and three transmembrane segments (Fig 2C). Members of the genus *Benyvirus* have between four or five linear positive sense ssRNAs capped at the 5′ end (Kiguchi et al., 1996; Saito et al., 1996); however, despite an exhaustive search, we failed to identify RNAs coding for movement proteins, or CP proteins. This fact is unsurprising as benyviruses can lose some genomic RNAs (Bouzoubaa et al., 1991; Heidel et al., 1997; Kondo et al., 2013; Lozano and Morales, 2009). The taxonomic status of these Beny-like sequences should be clarified by the ICTV.

**Figure 2.**
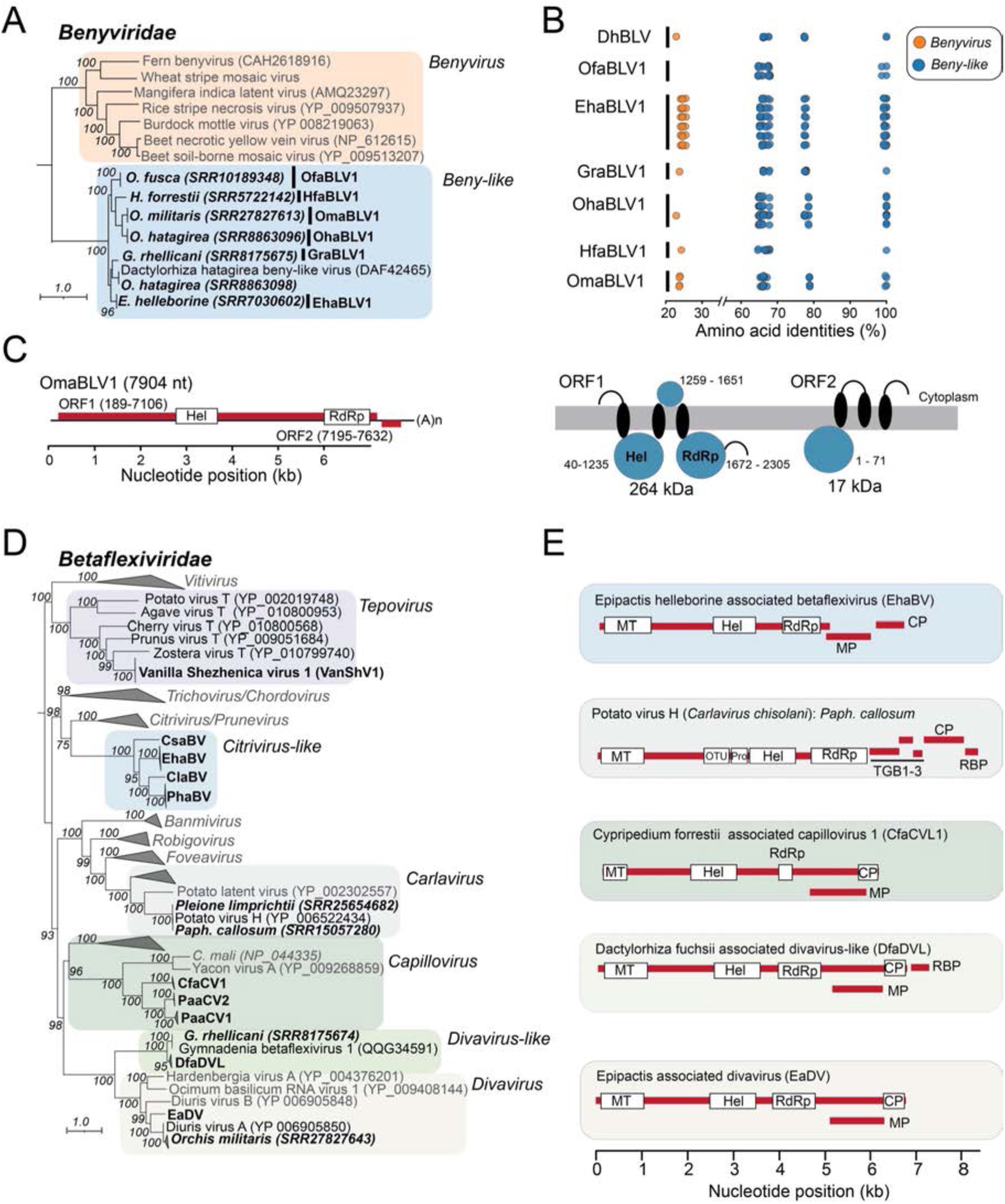
Description of novel viruses detected in RNA-seq data from orchids (Part I). A) The phylogenetic tree of Benyvirus-like sequences detected in the RNA-seq data from orchids suggests the existence of a new genus within *Benyiviridae.* B) comparison of the amino acid identities of the ORF1 protein of Beny-like viruses (blue) to members of the genus *Benyvirus* (orange). C) genome features typical of Beny-like viruses illustrate the position of predicted Hel and RdRp domains, along with a putative ORF2 encoding a 17 kDa protein (right). These proteins are expected to be embedded in a membrane, with the topology in the left panel. D) Phylogenetic tree illustrating the relationships among *Betaflexiviridae* associated with orchids. E) Genome structure of representative viruses within distinct lineages of *Betaflexiviridae*. Viruses detected in the RNA-seq data are shown in bold in the phylogenetic trees. For single sequences, the host and SRA accession codes are indicated. For multiple sequences, only the acronyms are provided.

### Betaflexiviridae

We identified several new virus-host associations, novel species, and possibly new genera within this family (Fig 2D). In the genus *Carlavirus*, we identified Potato virus H (PotVH, *Carlavirus chisolani*) sequences in *Paphiopedilum callosum* and *Pleione limprichtii* ∼98% identical to isolate YN from China (Li et al., 2013), representing the first report of PotVH in orchid species. In the genus *Tepovirus,* we found a putative new species in *Vanilla shenzhenica,* most closely related to zostera virus T (∼68% nt identities), which was independently identified by Choi et al. (2024) and named vanilla shenzhenica virus 1 (VanShV1) (Fig 2D). In the genus *Capillovirus*, we identified three putative new species associated with *Cypripedium forrestii* and *Paphiopedilum* species, which we named cypripedium forrestii associated capillovirus 1 (CfaCVL1), paphiopedilum-associated capillovirus 1 (PaaCV1), and paphiopedilum-associated capillovirus 2 (PaaCV2) (Fig 2D and E). These viruses exhibited the expected genomic features of capilloviruses (Fig 2E) and clustered in a clade sister to *C. mali* and yacon virus A (Liebenberg et al., 2012). In the genus Divavirus, we identified diuris virus A (DVA, *Divavirus alphadiuris*) sequences in *Dacthyloriza fuchsia*, *D. majalis*, and *O. militaris* and a new Divavirus in *Epipactis atrorubens* most similar to DVA (63% amino acid identities) which we named epipactis associated divavirus (EaDV) (Fig 2D and E).

We also identified a set of sequences in *Dactylorhiza fuchsia* most closely related to gymnadenia betaflexivirus 1 (GBFV1), which we named dactylorhiza fuchsii associated divavirus-like (DfaDVL). DfaDVL and GBFV1 formed a clade sister to the genus Divavirus and encoded an additional putative nucleic acid binding protein at the 5’ end (Wylie & Jones 2011) (Fig 2D and E). These sequences likely represent a novel genus within *Betaflexiviridae*. Finally, we found a clade comprising four putative new virus species associated with genera *Epipactis*, *Paphiopedilum*, *Cypripedium*, and *Collarorhiza,* which we named epipactis helleborine associated betaflexivirus (EhaBV, paphiopedilum henryanum associated betaflexivirus (PhaBV), cypripedium lichiangense associated betaflexivirus (ClaBV), and collarorhiza striate-associated betaflexivirus (CsaBV). These viruses clustered in a clade sister to the genera *Prunevirus* and *Citrivirus* and exhibited a genome structure similar to citriviruses (Vives et al., 2001). However, these citrivirus-like viruses had a smaller genome size (∼6.7 kb vs ∼8.7 kb), resulting from a smaller RdRp protein lacking AlkB and protease domains (Fig 2D).

### Bromoviridae

Based on GenBank records, CMV is the only virus from the family *Bromoviridae* known to infect orchid species. CMV has been found in *Cymbidium goeringii*, *Cymbidium* sp., *Vanilla* x *tahitensis*, *Calanthe discolor,* and *V. planifolia* (Madhubala et al., 2005; Wylie et al., 2015). We detected CMV in two additional species: *Cymbidium sinense* and *Dendrobium moniliforme* (Supp. Fig. 3D). These isolates are unrelated and likely originate from independent cross-transmission events, a common occurrence in this high generalist virus (Scholthof et al., 2011).

### Caulimoviridae

No viruses from this family have been reported in Orchidaceae. Here, we report seven tentative new *Badnavirus* species in *Mexipedium xerophyticum*, *Dendrobium catenatum*, and *Paphiopedilum armeniacum* (Fig. 3A). For these species, we propose the tentative names mexipedium_xerophyticum associated badnavirus 1 (MaBV1), dendrobium_catenatum associated badnavirus 1 (DaBV1), and paphiopedilum associated badnavirus 1-4 (PaBaV1-4). Analysis of the encoded proteins revealed that MaBV1 was most similar to *B. betamaculaflavicannae* (∼85% a.a. identities), DaBV1 to pelargonium vein banding virus (59.4%), PaBaV1 to *B. alphavirgamusae* (66.5%), and PaBaV2 to *B. etadioscoreae* (61.4%). PaBaV3, on the other hand, forms a distinct lineage distantly related to *B. decoloratiovitis* (58.4%). Lastly, PaBaV4 and PaBaV5 were most closely related to each other (73.1%) and formed a clade sister to *B. vitis*, *B. fici*, and *B. decoloratiovitis*.

**Figure 3.**
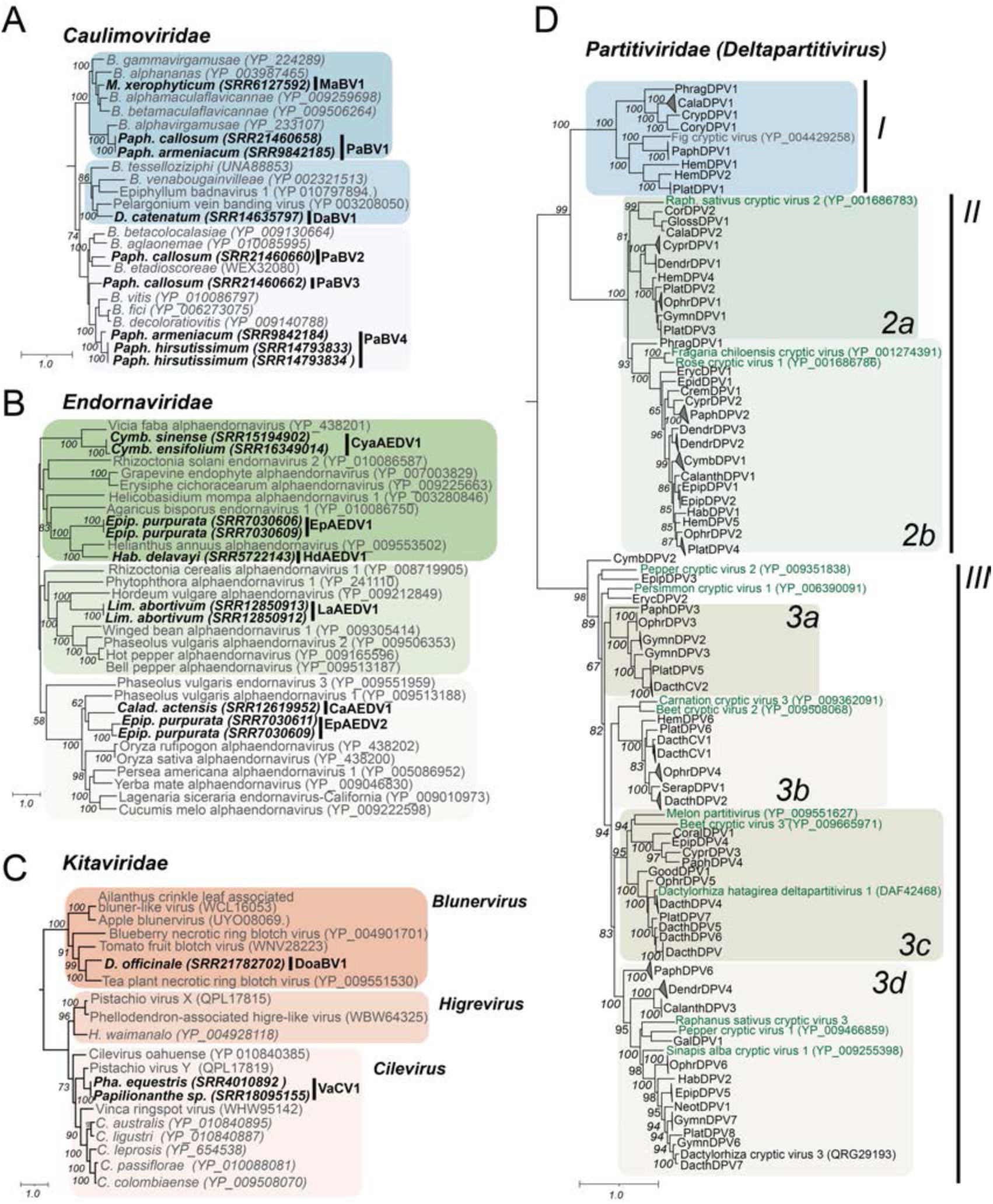
Description of novel viruses detected in RNA-seq data from orchids (Part II). Phylogenetic trees illustrating the phylogenetic relationship of viral sequences from the families *Caulimoviridae* (A), *Endornaviridae* (B), *Kitaviridae* (C), and *Partitiviridae* (D). Viruses identified in the RNA-seq data are shown in bold in the phylogenetic trees in panels A, B, and C and are labeled with the orchid host, SRA accession, and acronyms. In panel D, major clades are indicated using Roman numerals. Reference sequences are shown with full names in green. Sequences detected in RNA-seq data are labeled with their assigned acronyms; details are provided in the supplementary data.

### Endornaviridae

Our analysis revealed six new alphaendornaviruses in *C. sinense* and *Cymbidium ensifolium* (CyaAEV1), *Epipactis purpurata* (EpAEDV1 and EpAEDV2), *Habenaria delavayi* (HdaAEDV1), *Limodorum abortivum* (LaAEDV1), and *Caladenia actensis* (CaAEDV1) (Fig 3B). These viruses clustered into four clades: CyaAEV1, clustered with vicia faba alphaendornarvirus; EpAEDV1 and HdaAEDV1 formed a clade with helianthus annuus alphaendornavirus; LaAEDV1 was most closely related to hordeum vulgare alphaendornavirus, while CaAEDV1 and EpAEDV2 where more closely related to phaseolus vulgaris alphaendornavirus (Fig 3B).

### Geminiviridae

Begomovirus spathoglottis 1 (*Begomoviru*s) is the only *Geminiviridae* reported in Orchidaceae (GenBank: OQ791967, unpublished). We identified a partial sequence 85.5% identical to chickpea redleaf virus 2 (CpRLV2, *Mastrevirus*) in *Phalaenopsis equestris* (Filardo and Sharman 2019) and another 99.8% identical to tomato yellow leaf curl virus (TYLCV, *Begomovirus*) in *C. appendiculata* (Supp. Fig 3E). These findings are the first reports of CpRLV2 and TYLCV in orchids.

### Kitaviridae

Solanum violifolium ringspot virus (SvRSV, *Cilevirus. solani*) is the only known *Kitaviridae* in orchids (Ramos-González et al., 2022). We identified two new members of the *Kitaviridae* from the genera *Cilevirus* and *Blunervirus* (Fig. 3C). The Cilevirus was identified in *Papilionanthe* sp. and *Pha. equestris* and was most similar to pistachio virus Y (GenBank QPL17819, 61.3% amino acid identities). As *Papilionanthe* and *Phalaenopsis* are genera within the tribe Vandae, we propose the name vandae-associated cilevirus 1 (VaCV1) for this virus. The *Blunervirus* was detected in *Dendrobium officinale* and was most closely related to the tea plant necrotic ring blotch virus (43.4% amino acid identities, GenBank YP_009551530). For this virus, we propose the name dendrobium officinale associated blunervirus 1 (DoaBV1).

### Partitiviridae

We focused our analysis on the genus *Deltapartitivirus,* which is a genus that exclusively infects plants (Vainio et al., 2018). Four deltapartitivirus have been reported in orchids: dactylorhiza hatagirea deltapartitivirus (DacthDPV1), and dactylorhiza cryptic virus 1-3 (DacthCV1-3) (Sidhartan et al., 2021; Wylie et al., 2013a). We confirmed the presence of DacthCV1, DacthCV2, and DacthDPV1 in the datasets from *Dactylorhiza fuchsii* and *Dactylorhiza incarnata* (Figure 3D). In addition, we found 72 putative new *Deltapartitivirus* species, which we classified based on RdRp similarities (90%) and could be subdivided into three clades (Figure 3D). Clade I included sequences clustered with Fig cryptic virus (Fig 3D). Clade II comprised 26 sequences subdivided into two subclades based on their relatedness to raphanus sativus cryptic virus 2 (2a) and fragaria chiloensis cryptic virus (2b). Finally, clade III comprised sequences species subdivided into several subclades: 3a, related to DacthCV2; 3b, related to DacthCV1, carnation cryptic virus 3 and Beet cryptic virus 2; 3c, related to dactylorhiza hatagirea deltapartitivirus 1, melon partitivirus and beet cryptic virus 3; and 3d comprising sequences related to Raphanus sativus cryptic virus 3, pepper cryptic virus 1 and sinapsis alba cryptic virus 1. Deltapartitiviruses were more prevalent in Cypripedioideae and Orchidoideae and less prevalent in Epidendroideae (Supplementary Fig 2B).

### Phenuiviridae

Members of this family infect plants, animals, and fungi but have not been described in orchids (Sasaya et al., 2023). We identified four rubodvirus-like sequences in *Erycina crista-galli*, *Pleione limprichtii*, *Cypripedium lichiangense*, *C. fargesii*, and *C. forrestii* (Fig 4A, Supp. data). A BLASTx analysis of the encoded CP proteins revealed identities in the 30.0-42.9% range and protein similarities in the 50.6-58.2% range to reference species. The *Erycina crista*-*galli* sequences were identified in nine datasets and were most similar to Qingdao RNA virus 3, an uncharacterized virus detected in leaf tissue from unidentified plants in China (Cao et al., 2022) (Fig 4 A). We propose the name and erycina-associated phenuivirus-like 1 (EaPhV1) for this virus. For the remaining sequences, we propose the names pleione-associated phenuivirus-like 1 (PaPhV1), cypripedium-associated phenuivirus-like 1 (CaPhV1), and cypripedium-associated phenuivirus-like 2 (CaPhV2). EaPhV1, PaPhV1, CaPhV1, and CaPhV2 formed clades outside *Rudobvirus* and might represent new genera (Figure 4A).

**Figure 4.**
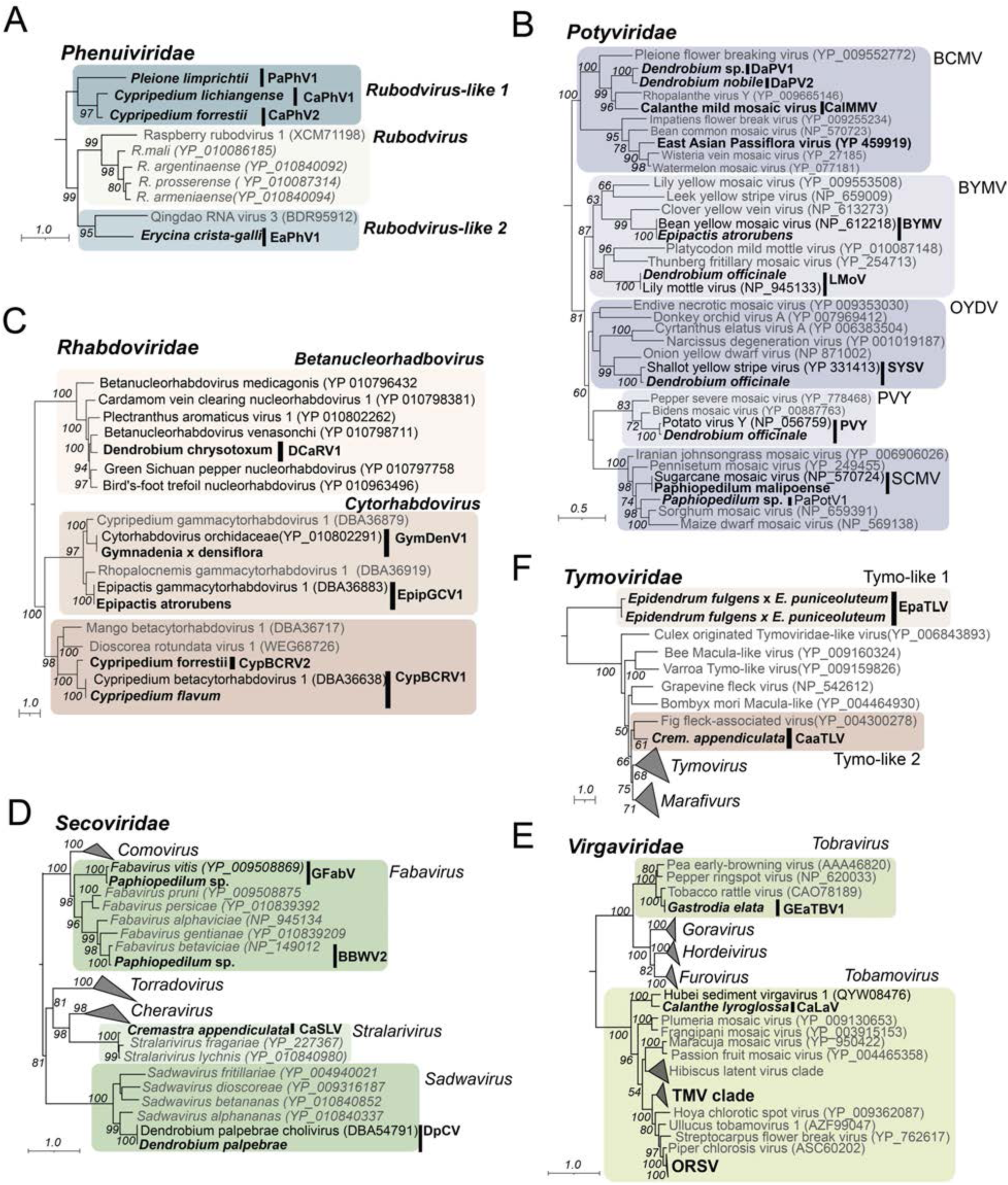
Description of novel viruses detected in RNA-seq data from orchids (Part III). Phylogenetic trees illustrating the phylogenetic relationship of viral sequences from the families *Phenuiviridae* (A), *Potyviridae* (B), *Rhabdoviridae* (C), *Secoviridae* (D), *Tymoviridae* (E), and *Virgaviridae* (F). Viruses identified in the RNA-seq data are highlighted in bold. In panel B, potyvirus lineages are shown outside each clade. Sequences found in the RNA-seq data are labeled with their respective acronyms; further details are available in the supplementary data.

### Potyviridae

We detected ten potyviruses from six lineages, including four tentative new species (Fig 4B). In the BCMV lineage, we found calanthe mild mosaic potyvirus (CalMMV*, Potyvirus callanthis*) sequences in *Calanthe hoshii* sharing 99.1% nucleotide identities with the reference sequence from Japan (GenBank AB011404) (Gara et al., 2008). Prior to this work, only partial sequences comprising the coding region for CP and a part of NIb were available, and these sequences will represent the first nearly complete CalMMV genomes. In this lineage, we also identified two new potyviruses in *Dendrobium* species, which we tentatively named dendrobium-associated potyvirus 1 (DaPV1) and 2 (DaPV2). These viruses formed a distinct clade alongside Rhopalante virus Y and CalMMV, suggesting the existence of a distinct orchid lineage of potyviruses (Fig 4B). In this group, we also identified a partial sequence of east asian passiflora virus (EAPV, *Potyvirus orionspassiflorae*) in *Paphiopedilum concolor* x *Paphiopedilum hirsutissimum* most similar (88.35%) to isolate CY from Taiwan (KX520664). This finding would represent the first report of EAPV in orchids. In the BYMV lineage, we detected BYMV (*Potyvirus phaseoluteum*) sequences in *Epipactis atrorubens* that were most similar (94.7%) to strain Gla from Japan (Nakazono-Nagaoka et al., 2009). Additionally, we found several sequences of lily mottle virus (LMoV, *Potyvirus lilimaculae*) in *Dendrobium officinale*, sharing about 98.6% nucleotide identity with an isolate from tulip bulbs (van Gent-Pelzer et al., 2024) (Fig 4B). In the OYDV lineage, we identified several shallot yellow stripe virus (SYSV, *Potyvirus ascaloniae*) sequences in *D. officinale* closely related to isolate LL01 from Tiger lily (*Lilium lancifolium*, 97.8%). In the PVY lineage, we identified PVY in *D. officinale* most similar to potato isolates from *S. tuberosum* from China (99.7%) NMG-1 (MN607709) and TC (MF033142). In the SCMV lineage, we detected sugarcane mosaic virus (SCMV, *Potyvirus sacchari*) in *Paphiopedilum malipoense* (Chen et al., 2002). In this group, we also found a putative new potyvirus species in the hybrid *Paphiopedilum concolor* x *Paphiopedilum hirsutissimum,* which we named Paphiopedilum-associated potyvirus 1 (PaPotV1), more related to Sorghum mosaic virus (SrMV, 72 % nucleotide identities) (Fig 4B). Finally, in the TEV lineage, we found a 988 nt contig from *Paphiopedilum concolor* x *Paphiopedilum hirsutissimum* sharing 66.23% identities in the CP region of Tobacco etch virus (TEV) which suggests the existence of new potyvirus which we named Paphiopedilum-associated potyvirus 2 (PaPotV2) (Supp. Fig 4C).

### Rhabdoviridae

In this family, we identified sequences associated with the genera *Betanucleorhaddovirus, Cytorhabdovirus,* and *Varicosavirus* (Fig. 4C, Supplementary Fig. 4D-F). In the genus *Betanucleorhabdovirus*, we discovered a putative new virus in *Dendrobium_chrysotoxum* related to *Betanucleorhabdovirus venasonchi*, which we named dendrobium chrysotoxum-associated rhabdovirus 1 (DCaRV1). Findings in the genus *Cytorhabodvirus* were more numerous.

We identified cypripedium betacytorhabdovirus 1 (CypBCRV1) in *Cypripedium flavum* (Bejerman et al., 2021, 2023; Dietzgen et al., 2014) and a putative new species in *C. forrestii,* which we named cypripedium betacytorhabdovirus 2 (CypBCRV2). We also identified gymnadenia densiflora virus 1 (GymDenV1, *Cytorhabdovirus orchidaceae*) in *Gymnadenia_x_densiflora* and epipactis gammacytorhabdovirus 1 (EpipGCV1) in *Epipactis atrorubens* (Bejerman et al., 2023; Bejerman et al., 2021). Further analysis of partial sequences revealed the presence of *Cytorhabdovirus hordei* (BYSMV) in *Dendrobium officinale,* a putative new gammacytorhabdovirus in *Platanthera guangdongensis* (Platanthera gammacytorhabdovirus-like 1, PlaGCRV1), most closely related to coptys gammacytorhabdovirus 1 (N protein DBA36860, 47% aa identities) and a partial CP sequence that likely represents a novel varicosavirus in *Orchis italica* (OVL1(Orchis italica associated varicosavirus-like 1, OIaVL1) with 51% aa identities with Xinjiang varicose-like virus (GenBank YP_010840821) (Supp. Fig. 4C-D).

### Secoviridae

Three *secoviridae* species have been reported in orchids: canberra spider orchid nepovirus, habenaria delavayi nepovirus, and dendrobium palpebrae choliviru*s* (Sidhartan et al., 2024). In the genus *Fabavirus* we identified grapevine fabavirus (GFabV, *Fabavirus vitis*) in *Paphiopedilum concolor* x *Paphiopedilum hirsutissimum*, and broad bean wilt virus 2 (BBWV2, *Fabavirus betaviciae*) in *Paphiopedilum concolor* x *Paphiopedilum hirsutissimum* and *Dendrobium officinale* (Figure 4D). In the genus *Sadwavirus* we found dendrobium palpebrae cholivirus (DpCV) in *Dendrobium palpebrae*. We also found a new virus in the genus *Stralavirus*, which we named cremastra appendiculata stralavirus (CaSLV). This virus shared 78.0%-83.2% with camellia sinensis secoviridae 1 and 74.7%-82.5% with lychnis mottlevirus (*Stralarivirus lychnis*). We also found three partial sequences between 596-932 nt in length sharing 80.7-83.7% nucleotide identities with Cnidium vein yellowing virus 2 in *Cymbidium goeringii,* which likely represents a new isolate of this virus (Park YC et al., 2023).

### Solemoviridae

Three members of this family have been reported in orchids: cymbidium chlorotic mosaic virus (*Sobemovirus*), cymbidium chlorotic spot virus (*Sobemovirus*), and pterostylis polerovirus (*Polerovirus*). We identified partial sequences that likely represent a new *Polerovirus* species in *Dendrobium williamsonii*, related to polerovirus monocotyledonae 2 (< 83% amino acid identities) and pterostylis polerovirus (< 65.5 % amino acid identities) (Supp. Fig 4A). We propose the name dendrobium williamsonii-associated polerovirus 1 (DWaPV1) for this virus.

### Tombusviridae

The only *Tombusviridae* known to infect orchids is carnation mottle virus (CarMV, *Alphacarmovirus dianthi*), reported in *Phalaenopsis* hosts (Zheng et al., 2011b). This virus was identified as a partial sequence of 1,231 nt in *Dendrobium nobile* (SRR18516560) 99.1% identical to isolate DSMZ PV-1234 (OM867874) (Supplementary Fig. 4B)

### Tospoviridae

capsicum chlorosis virus (CaCV*, Orthotospovirus capsiciflavi*) in *Oncidium* sp. was the only *Tospoviridae* detected in this study. CaCV has already been reported in several orchidaceous hosts (Zheng et al., 2011a).

### Tymoviridae

No members of the *Tymoviridae* have been previously reported in Orchidaceae. We found two tymoviridae-like viruses in *Epidendrum puniceoluteum* (Fig. 4F), which we tentatively named epidendrum puniceoluteum-associated tymo-like virus (EpaTLV) and cremastra appendiculata-associated tymo-like virus (CaaTLV). At the amino acid level, EpaTLV was most closely related to tomato blistering mosaic virus and bee Macula-like virus (27%) and CaaTLV to glehnia littoralis marafivirus (∼77%).

### Virgaviridae

Unsurprisingly, the most common *Virgaviridae* was ORSV, from which we identified 89 samples from about a dozen species and five genera from the tribes Cymbidieae, Gastrodieae, Vandeae, Epidendreae, and Cranichideae in the subfamilies Orchidoideae and Epidendroideae (Supplementary Table). We also identified a putative new *Tobravirus* in *Gastrodia elata*, which we tentatively named gastrodia elata-associated tobravirus (GEaTBV1). GEaTBV1 shared 73.9% nucleotide identities and 78.9% amino acid identities with tobacco rattle virus (TRV, *Tobravirus tabaci*) (Fig 4E). We also identified a nearly complete genome of a new tobamovirus in *Calanthe lyroglossa*, most closely related to hubei sediment virgavirus 1 (78% amino acid identities, 71.7% nucleotide identities, 18% coverage), which was identified in a metagenome analysis of viruses in China (Chen et al., 2022). We propose the tentative name calanthe lyroglossa-associated virgaviridae (CaLaV) for this virus. Finally, we found a hordeivirus-like sequence of 477 nt in Gymnadenia rhellicani sharing 58% identity with the gamma B protein of Lychnis ringspot virus (YP_009508259), which we named gymnadenia rhellicani associated hordeivirus (GraHV) (Supp. Fig 4G).

### Virus associations

We then examined the co-occurrence of viruses within the same dataset. Except for *Solemoviridae, Tombusviridae,* and *Tospoviridae*, at least one member of each virus family was found in association with another (Fig. 5A). *Partitiviridae* was the family associated with the largest number of viruses, being found together with members of fifteen other families (Fig. 5A). This fact is not surprising considering the cryptic and persistent nature of these viruses and their high prevalence across datasets (Fig. 1A). *Amalgaviridae* and *Endornaviridae*, the other cryptic viruses in this study, were only found alongside *Partitiviridae* (Fig 5A). *Alphaflexiviridae* was the second family most frequently associated with other viruses. It was identified along with seven virus families with a remarkably strong association with *Virgaviridae* due to the co-occurrence of ORSV and CymMV in 60 samples. The third most common coinfections involved *Betaflexiviridae*, which, besides *Partitiviridae* and *Alphaflexivirida*e, was also found together with members of *Rhabdoviridae*, *Secoviridae*, *Phenuviridae*, and *Benyviridae* (Fig 5A). We then examined the enrichment or depletion of one virus family in the presence of another using Fisher’s exact test (Fig 5B). We found fourteen positive and six negative correlations between virus families. The most significant positive associations were found between the pairs *Alphaflexiviridae*/*Virgaviridae* (*P*<6.5×10^-29^), *Rhabdoviridae/Partitiviridae* (*P*<.0007), and *Partitiviridae*/*Secoviridae* (*P*<2.5×10^-7^) (Fig 5A-B). Interestingly, we found a negative association between *Alphaflexiviridae*/*Partitiviridae* (*P*<1.64×10^-13^); *Virgaviridae/Partitiviridae* (*P*<1.9×10^-5^); *Rhabdoviridae*/*Alphaflexiviridae* (*P*<3.2×10^-7^); *Betaflexiviridae/Alphaflexiviridae* (*P*<8.9×10^-5^); and *Secoviridae*/*Alphaflexiviridae* (*P*<.001).

**Figure 5.**
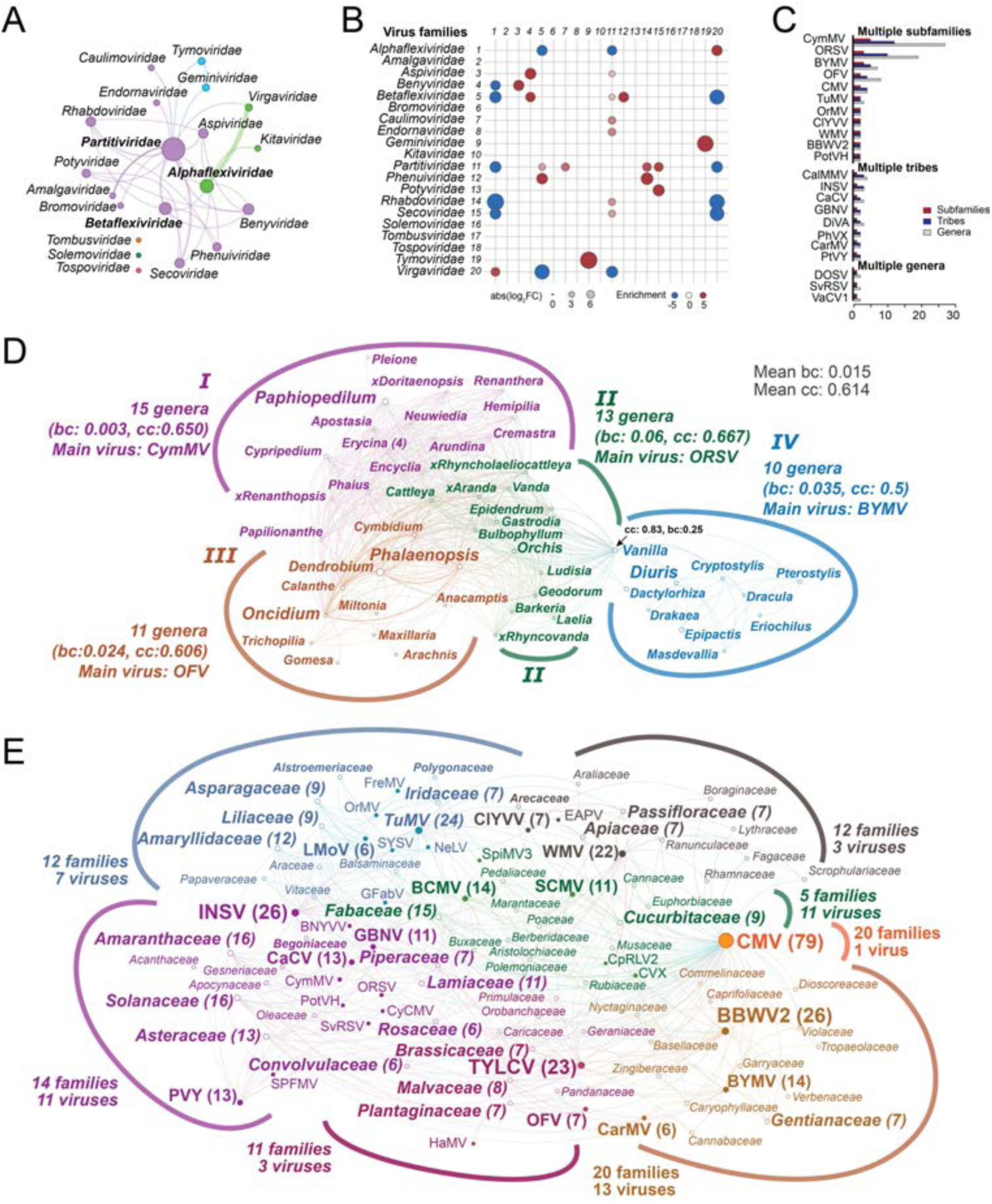
Biological associations of orchid viruses. A) Monopartite network illustrating the co-occurrence of virus families within the same dataset. Edge thickness is proportional to the number of co-occurrences. B) Correlation plot illustrating virus families significantly enriched (red) or depleted (blue) relative to the viruses displayed on the y-axis. The size of the circles represents the fold enrichment on a log2 scale. C) Bar plots illustrating the number of orchid taxa associated with generalist orchid viruses. D) A monopartite network illustrating the generalist viruses shared among orchid genera. Edges represent individual viruses. Colors represent network communities. In each instance, the average betweenness centrality (bc), closeness centrality (cc), and dominant virus are specified. E) Bipartite network illustrating the association of orchid viruses (closed circles) with other plant families (open circles). Colors indicate distinct communities within the network. The number of associations for the most significant nodes is indicated in parentheses. Panels C, D, and E also include orchid viruses reported in GenBank but not found in this study. Detailed information about the networks is provided in the supplementary material.

### Host range of orchid viruses

To gain further insights into the host range of orchid viruses across Orchidaceae, we combined our findings with all the viruses reported to infect orchids in the NCBI virus database. According to this, the current orchid virome includes about 213 viruses across 70 orchid genera, with an average of 3.4 virus species linked to each genus (Supp. Fig 2A). Orchid viruses were very specific, with 191 (89.9%) viruses limited to a single genus and only 22 viruses affecting more than two genera (Figure 5C). The most generalist viruses were CymMV and ORSV, which infected species from 27 and 19 genera, respectively. Other viruses capable of infecting multiple subfamilies include BYMV, OFV, CMV, TuMV, OrMV, WMV, BBWV2, and PotVH. Eight viruses were identified as restricted to a single orchid subfamily yet capable of infecting multiple tribes: CalMMV, INSV, CaCV, GBNV, DiVA, PhVX, CarMV, and PtVY. Finally, three viruses were limited to different genera within the same tribe: DOSV and SvSRV were limited to Orchidoideae, and VaCV1 to Epidendroideae. The genera with the highest number of virus species were *Dendrobium* (31 viruses), *Paphiopedilum* (22 viruses), *Gymnadenia* (17 viruses), and *Phalaenopsis* (14 viruses).

We then used network analysis to understand the association of generalist viruses within Orchidaceae (Fig 5D). The network depicted orchid genera as nodes and the viruses shared between genera as edges. Generalist viruses formed a giant component comprising 49 orchid genera that shared between 1 and 6 viruses. The genera with the largest number of shared viruses were the pairs *Calanthe/Dendrobium* (6 viruses), *Calanthe/Cymbidium* (5), *Phalaenopsis/Oncidium* (5), *Phalaenopsis/Dendrobium* (5), and *Cymbidium/Dendrobium* (5). This network can be subdivided into four communities. Community I comprised fifteen orchid genera, all of which were connected to the genus *Paphiopedilum,* which acted as a hub; CymMV was the most abundant virus in this subnetwork (Fig. 5D). Community II comprised thirteen genera and was highly connected, as all nodes were linked to each other; the most common virus was ORSV. Community III comprised eleven genera, with *Phalaenopsis* and *Oncidium* as hubs (nine connections each); in this subnetwork, OFV was the most frequent virus. Finally, community IV comprised ten viruses, with *Diuris* acting as a hub with eight connections; BYMV was the most frequent virus in this community. Closeness centrality measures how close a node is to all others in the network. Globally, the network exhibited a mean closeness centrality of 0.614. Community IV exhibited closeness centrality below the average 0.5, indicating that these orchid genera share fewer viruses in common with the remainder of the network. Another informative network parameter is betweenness centrality, which measures the importance of a node in connecting other nodes. The network exhibited a mean betweenness centrality of 0.015. Interestingly, the genus Vanilla displayed a significantly higher value of 0.255 and possessed the highest closeness centrality, indicating its role in linking the virome of community IV to the rest of the network (Fig. 5D).

### Alternative hosts

33 (15.5%) orchid viruses were found to infect plant species from other families. Orchidaceae has the largest number of viruses in common with species from Amaranthaceae (16), Solanaceae (16), Fabaceae (15), Asteraceae (13), Amaryllidaceae (12), and Lamiaceae (11). We examined the interaction between alternative hosts using a bipartite network where nodes represent plant families and virus species, and edges indicate virus-host associations (Fig 5F). Viruses exhibited a mean closeness centrality of 0.387 and a mean betweenness centrality of 0.006. Similar values were observed for plant families, which exhibited mean closeness and betweenness centralities of 0.335 and 0.0034, respectively. Interquartile range analysis revealed that CMV is an outlier for closeness centrality (0.644), while CMV (0.595), TuMV (0.073), INSV (0.065), and WMV (0.064) were identified as upper bound outliers in betweenness centrality. These viruses are well-known generalists and were found to infect more than 20 different families (Fig 5F). A similar analysis of plant nodes revealed that Amaranthaceae (0.480), Solanaceae (0.480), Fabaceae (0.480), Asteraceae (0.469), Lamiaceae (0.463), and Cucurbitaceae (0.453) are upper bound outliers in closeness centrality. Additionally, eleven plant families were identified as upper outliers in betweenness centrality: Amaranthaceae (0.085), Solanaceae (0.076), Fabaceae (0.063), Amaryllidaceae (0.034), Asteraceae (0.031), Asparagaceae (0.029), Plantaginaceae (0.026), Lamiaceae (0.026), Liliaceae (0.024), Passifloraceae (0.022), Malvaceae (0.017), Cucurbitaceae (0.016), Polemoniaceae (0.015), and Iridaceae (0.014).

The alternative host network can be divided into six communities or subnetworks, which we named after the nodes with the highest local closeness centrality: *Solanaceae, Fabaceae*, CMV, TuMV, BBWV2, WMV, and TYLCV. In the Solanaceae and Fabaceae subnetworks, a plant host acted as a central node, indicating a significant role as an alternative host of orchid viruses. The Solanaceae subnetwork comprised fourteen plant families and eleven viruses, among which the most prominent were INSV and CaCV. It also included major alternative hosts such as *Amaranthaceae*, *Asteraceae*, and *Lamiaceae*. The Fabaceae subnetwork included five viruses and 16 families. The dominant viruses were BCMV and SCMV, including major hosts Cucurbitaceae and Polemoniaceae. The remaining subnetworks represented communities where viruses acted as central nodes. The largest is the CMV community, which includes 20 families with CMV, the only virus shared with Orchidaceae. The TuMV community included six additional viruses (LMoV, OrMV, NeLV, FreMV, SYSV, GFabV, and families, of which the most important were Amarylidaceae, Asparagaceae, Liliaceaee, and Iridaceae (0.014). The BBWV2 community included two additional viruses (CarMV and BYMV) and thirteen families; none were considered primary alternative hosts. The WMV subnetwork also included ClYVV, EAPV, and 12 plant families. The most notable family is Passifloraceae. Finally, the TYLCV also included OFV and HaMV and 11 families, of which the most notable were Plantagiancea and Malvaceae (2 viruses)

### Transmission mechanisms

A cluster analysis of transmission mechanisms for the virus genera reported to infect orchids revealed nine main groups (Fig. 6A). The largest included genera efficiently transmitted through sap and vegetative propagation. This group can be further subdivided into seed-borne (*Goravirus, Tepovirus, Citrivirus, Nepovirus, Sadwavirus,* and *Capillovirus*), aphid transmissible (*Potyvirus, Carlavirus, Fabavirus*), and genera poorly transmitted by aphid or seeds (*Hordeivirus, Tymovirus, Sobemovirus, Potexvirus, Platypuvirus, Tobamobirus, Alphacarmovirus, Cytorhabdovirus*). The second cluster in size comprised genera efficiently transmitted through vegetative propagation but not through sap, which could be subdivided into mite-transmitted (*Dichorhavirus, Cilevirus,* and *Allexivirus*) and non-mite transmitted (*Badnavirus, Divavirus,* and *Amalgavirus*). The remaining clusters involved viruses more efficiently transmitted through seeds (*Alphaendornavirus, Stralavirus, and Tobravirus*), aphids (*Poacevirus, Polerovirus, Betanucleorhabdovirus, Cucumovirus*), whiteflies (Begomovirus), thrips (*Orthotospovirus*), soil-borne viruses (*Blunervirus, Varicosavirus* and *Ophiovirus*), plasmidiophorids (*Benyvirus*), and leafhoppers (*Mastrevirus*). An analysis of cytochrome oxidase I (*CoxI)* barcodes in the RNA-seq data revealed the recurrent presence of insects from the families Aphididae (32), Thripidae (23), in addition to pollinating insects from the families Nymphalidae (48), Noctuidae (21), Bombycidae (20), and Apidae (12), that may act as carrier of pollen transmitted viruses (Bhat and Rao, 2020; Fetters et al., 2022,2023) (Supp. Fig 1F). Interestingly, a cluster analysis between virus genera and the lifestyle of orchids suggests differences between the viromes of epiphytic and terrestrial species (Fig 6B). This correlation between the ecology of hosts and the associated viruses could explain the correlation between viruses described previously.

**Figure 6.**
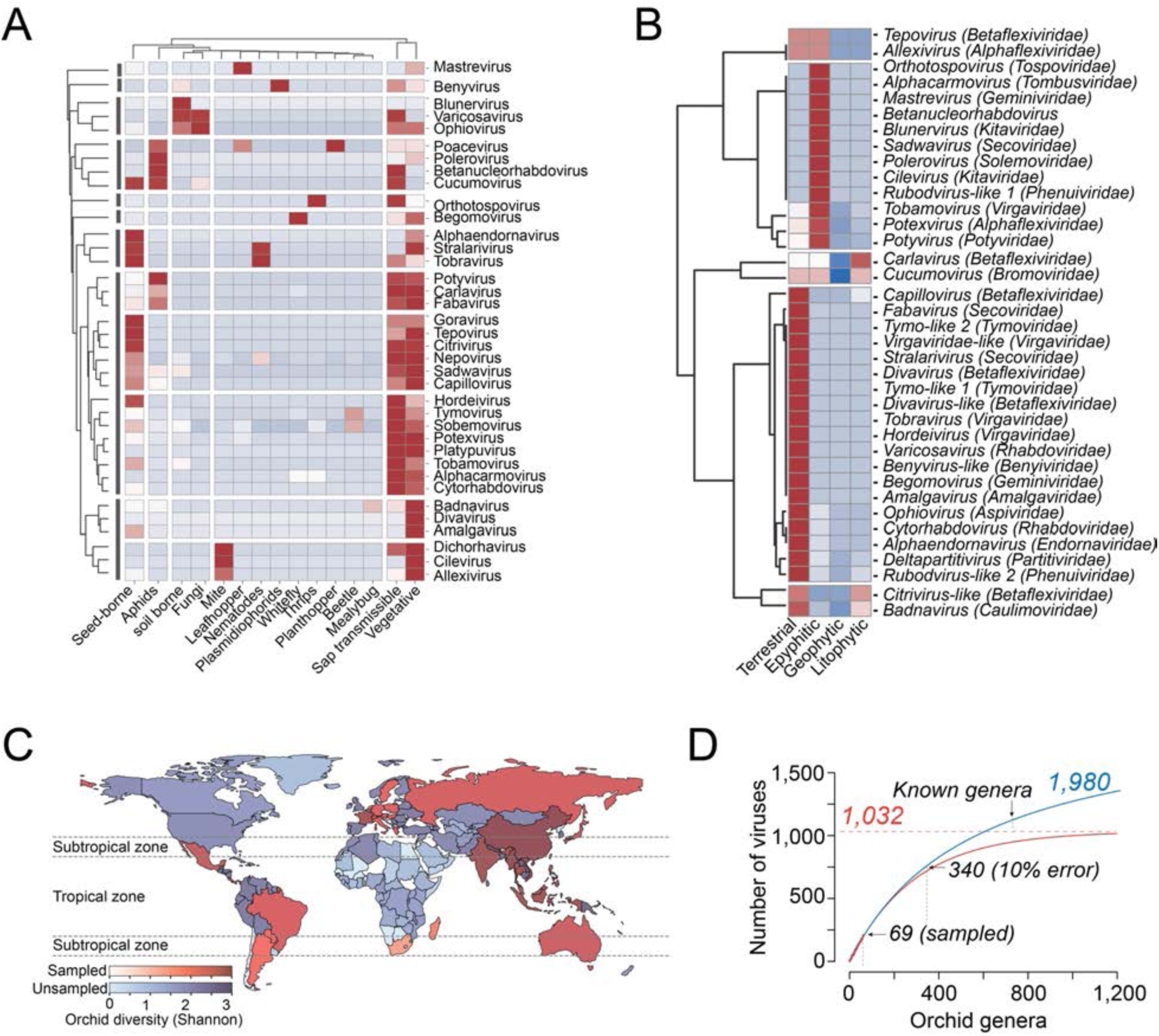
Ecological associations and virus diversity of orchid viruses. A) cluster map illustrating the transmission mechanisms of virus genera associated with orchids. The color scale represents the number of viruses within each genus known over (red) or underrepresented (blue) by the corresponding transmission mechanism. B) Cluster map illustrating the relationship between the habitats of orchid species and associated genera. Color values represent over (red) and under-represented (blue) virus genera. C) World map illustrating the orchid diversity of countries for which RNA-seq data has been collected (scale of red) or not (scale of blue). Color intensity is proportional to the Shannon diversity index. D) Rarefaction plots indicating the estimated virus diversity using negative exponential (red) and hyperbolic models (blue). Numbers indicate the estimated number of viral species estimated under each model.

### The size of the orchid virome

Orchidaceae is currently divided into more than 800 genera, but RNA-seq data is only available from approximately 40 of them; in addition, diversity hotspots from the Neotropics and Africa are missing or underrepresented (Fig. 6C). Considering these limitations, a rarefaction analysis indicates that the size of the Orchidaceae virome ranges from 1,032 (negative exponential model) to 1,980 (hyperbolic model) virus species. This preliminary estimate should be viewed with caution, as only viruses from 69 genera have been recorded, corresponding to the linear section of the rarefaction curve, with an estimated error of 1% at 120 genera and 10% at 340 genera (Fig. 6D). It is important to note that estimates of virus diversity may become unstable as discovery efforts intensify, as observed in studies on mammalian viruses where discovery rates are highly variable, with little evidence of declining towards an asymptote (Carlson et al., 2019; Gibb et al., 2022).

## DISCUSSION

Plant viruses pose one of the most significant threats to agriculture, and their impact is anticipated to grow due to global trade, human travel, and climate change (Das et al., 2021; Jeger et al., 2023; Jones, 2009; Ristaino et al. 2021; Singh et al. 2023). This menace is more pronounced in orchids, as many species are at high risk of extinction (Fay, 2018; Swarts & Dixon, 2009; Vitt et al., 2023). Our study confirms that we are still far from a complete understanding of the Orchidaceae virome, which is not surprising considering the high diversity, complex evolutionary history, and biogeography of this family (Givnish et al., 2015; Smith et al., 2018; Vitt et al., 2023). Unfortunately, our lack of knowledge about orchid viruses hinders the design of reliable virus detection tools and the implementation of knowledge-based integrated management strategies (Wylie et al., 2013). From a practical standpoint, the data presented offer a global perspective on the Orchidaceae virome that could aid in guiding future research and virus vigilance programs (Baldi and La Porta, 2020; Kalimuthu et al., 2022; Kanapiya et al., 2024; Mehetre et al., 2021). Generalist viruses are particularly interesting because they can potentially infect species across various genera, tribes, or even subfamilies and may contaminate germplasm collections and parent stocks used in orchid propagation (Das et al., 2021; Merrit et al., 2014; Popova et al., 2016). This issue is especially worrisome given that most viruses associated with orchids belong to genera that can be effectively transmitted through vegetative propagation (Fig 6C). Our study also confirms the high prevalence of CymMV and ORSV, reinforcing their status as the most important viruses in orchids (Chen et al., 2019; Koh et al., 2014; Ryu et al., 1995; Wong et al., 1997). These viruses were also the most widespread among countries, with CymMV found in datasets from nine countries and ORSV detected in four (Supplementary data). In addition, attention should be paid to BYMV and OFV, which have a worldwide distribution affecting a wide range of orchid genera and other economically important crops (Otero-Colina et al., 2021). We also have shown new associations with BBWV2, BYSM, EAPV, GFabV, LMoV, PotVH, PVY, SCMV, SYSV, and TYLCV, which are often associated with ornamentals in the families Amaryllidaceae and Liliaceae, as well as food crops in Fabaceae, Solanaceae, Passifloraceae, and Poaceae (Li et al., 2013; Gray et al., 2018; Prasad et al., 2020; van der Vlugt et al., 1999). These plant families are also cultivated in vast expanses of land, increasing the risk of cross-transmission due to high insect pressure and the expansion of the agricultural interface (Elena et al., 2014; Jones et al., 2009; Lefeuvre et al., 2019). However, we recognize that the significance of viral infections can only be assessed through careful experimentation, as not all viruses adversely affect their host (Rossinck, 2015). Viral infections can even have positive effects; for instance, mild strains can offer cross-protection against more severe strains, and for ornamental plants, they can enhance their beauty (Roossinck, 2011; Valverde et al., 2012).

It would also be interesting to investigate the strong association between cryptic viruses and orchids. Numerous examples of mutualistic relationships between cryptic viruses and their hosts have been discovered, including their ability to confer tolerance to drought, cold, and high soil temperatures, as well as to adapt to varying nutritional environments (Roossinck, 2011, 2015). This may apply to endornaviruses and partitiviruses, which can enhance plant tolerance to abiotic stresses but also disrupt RNA silencing and prime the immune system (Fukuhara et al., 2020; Horiuchi et al., 2003; Moriyama et al., 1996; Moriyama et al., 1999; Okada et al., 2011; Valverde and Gutierrez, 2007). This line of research will clarify the positive correlation of deltapartitiviruses with *Betaflexiviridae* and *Phenuiviridae*, as well as their negative correlation with *Alphaflexiviridae* and *Virgaviridae*. The latter interaction is of great interest because these families include CymMV and ORSV, the most significant orchid viruses, and may suggest biological control alternatives. Future work should also address the fundamental biology and molecular features of the viruses discussed here, especially the putative new genera within *Benyviridae*, *Betaflexiviridae*, *Phenuiviridae*, and *Tymoviridae*. Of particular interest are the biological transmission mechanisms of the Beny-like viruses, which, similarly to Benyviruses, might be transmitted by root-infecting vectors in the Plasmodiphorales (Gilmer et al., 2017; Rush, 2003). This would be consistent with the terrestrial habitats of their putative hosts *Epipactis helleborine*, *Gymnadenia rhellicani*, *Hemipilia forrestii*, *Orchis hatagirea*, and *Orchis militaris* (Pérez-Escobar et al., 2024; Zhang et al., 2018). Molecular studies should also investigate the role of the predicted ∼17 kDa polytopic membrane protein of ∼17 kDa and the unique features of the encoded RdRp protein lacking Methyltransferase and protease domains. Similar basic research should also clarify the basic biology of Citrivirus-like and Divavirus-like viruses and their taxonomic status as putative new genera within *Betaflexiviridae*. At the molecular level, studies should clarify the functioning of the smaller RdRp protein lacking AlkB and protease domains and confirm the role of the RBP protein in Divavirus-like viruses. We have also identified the first associations of Phenuiviridae with Orchidaceae, featuring viruses that belong to two Rudobvirus-like genera. Due to the limited knowledge about this virus family, future research should uncover unknown aspects, such as virion morphology and transmission mechanisms, which remain uncertain (Sasaya et al., 2023). Finally, the putative new genera in Tymoviridae, which tend to be highly transmissible viruses by sap and vegetative propagation and can be transmitted by beetles, should be confirmed and studied further (Dreher et al., 2012).

We hope this work shows how data mining can provide a cost-effective approach to gaining a global perspective on plant viruses. Global science often faces limitations due to insufficient resources, bureaucracy, regional laws, and local politics (Buckee, 2020; Carroll et al., 2018). However, data regional data, often gathered for non-epidemiological purposes, can be combined to reveal trends that are useful for the epidemiological surveillance of plant viruses (Park et al., 2018; Salathé et al., 2012). This strategy could be particularly advantageous in developing countries, where conducting detailed surveys using advanced technology can be difficult. Meta-analyses can also be used to justify global funding initiatives for research that lacks local support.

Finally, we emphasize the significance of open and accessible data to close gaps in generating knowledge and making informed, knowledge-based decisions (Tarkoma et al., 2020). We hope that the data provided will pave the way for new research directions and encourage future studies on the Orchidaceae virome, thereby deepening our understanding of plant viruses.

## Acknowledgments

We thank everyone involved in generating the data used in this study. This work was supported by Universidad Nacional de Colombia sede Medellín under Hermes Grant 61273.

---

**Supp. Fig.1.**
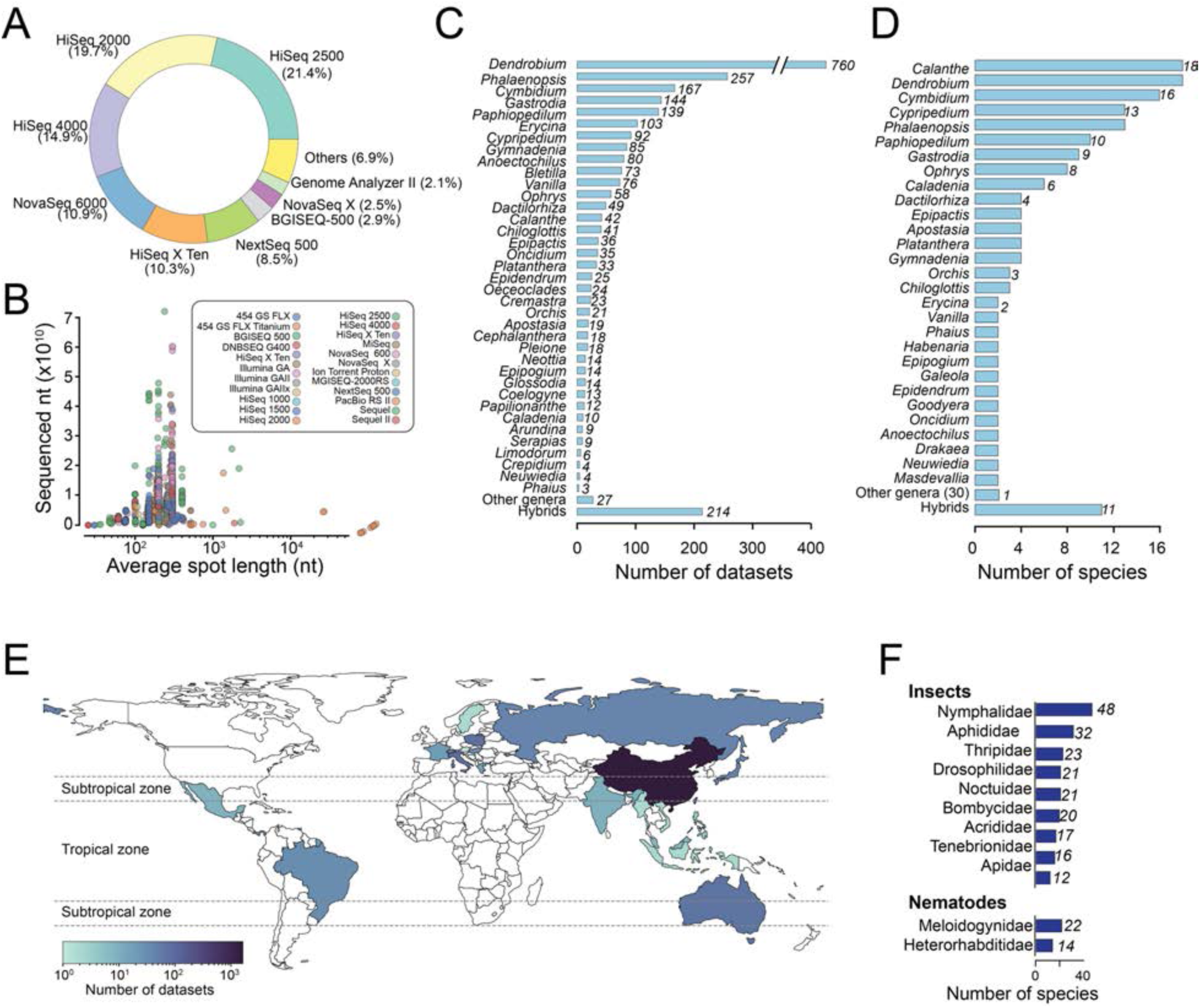
Global description of the datasets and findings. A) Donut chart illustrating the sequencing instruments used in the orchid datasets. B) Distribution of read lengths and sequenced depth in the datasets used in this study. C) Bar plot illustrating the number of datasets by orchid genera (D) and the number of orchid species per genera analyzed in this study. E) World map illustrating the countries of origin of the RNA-seq data used in this work. F) Bar plots depicting the most common insect and nematode families associated with orchid datasets based on *cox1* sequences.

**Supplementary Figure 2.**
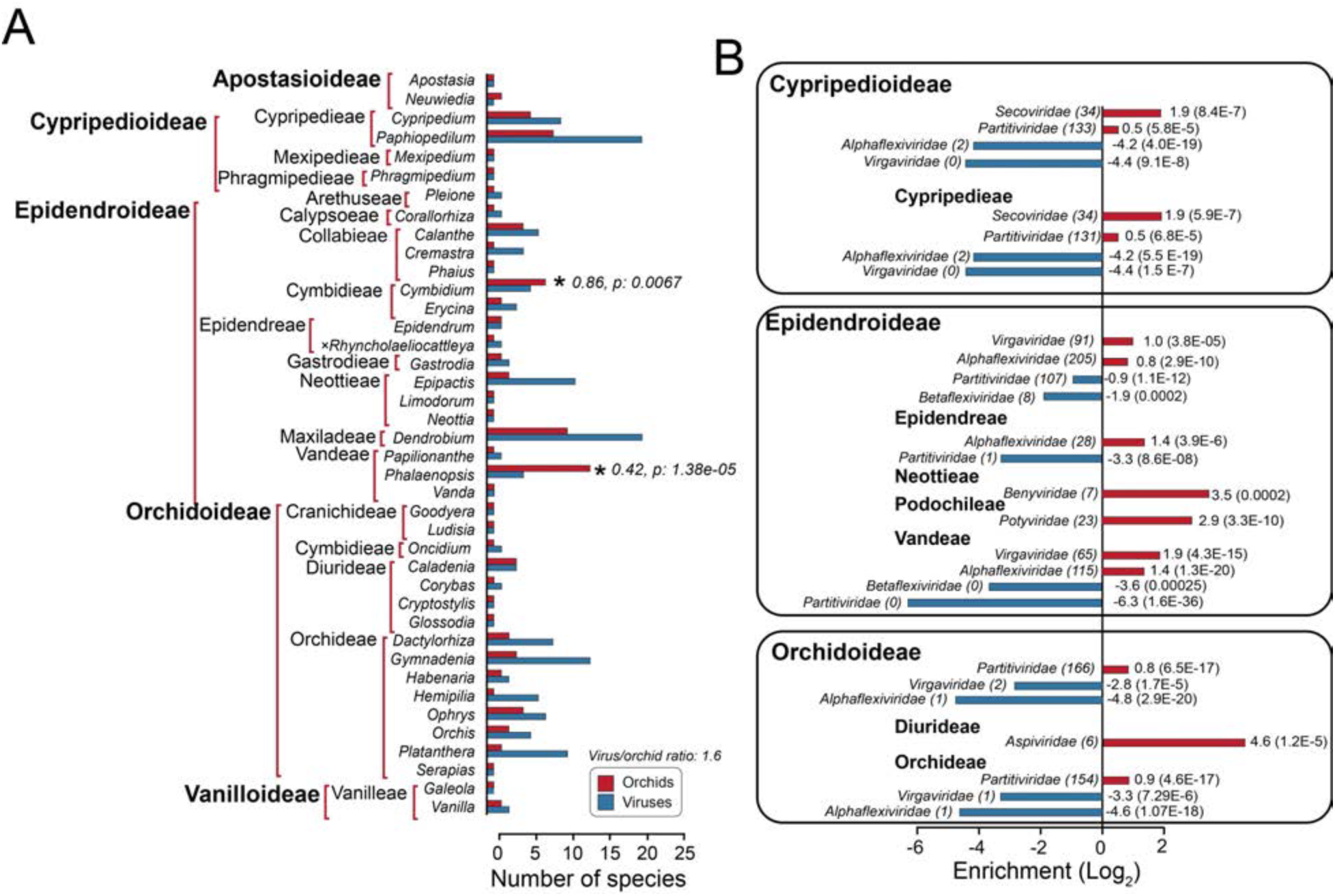
General relationships between viruses and orchid hosts. A) Bar plots illustrating the total number of orchid (red) and virus (blue) species detected in each genus. B) Virus families significantly enriched (red) or depleted (blue) in different orchid taxonomic ranks. P-values from Fisheŕs exact test are shown in parentheses.

**Supp. Fig.3.**
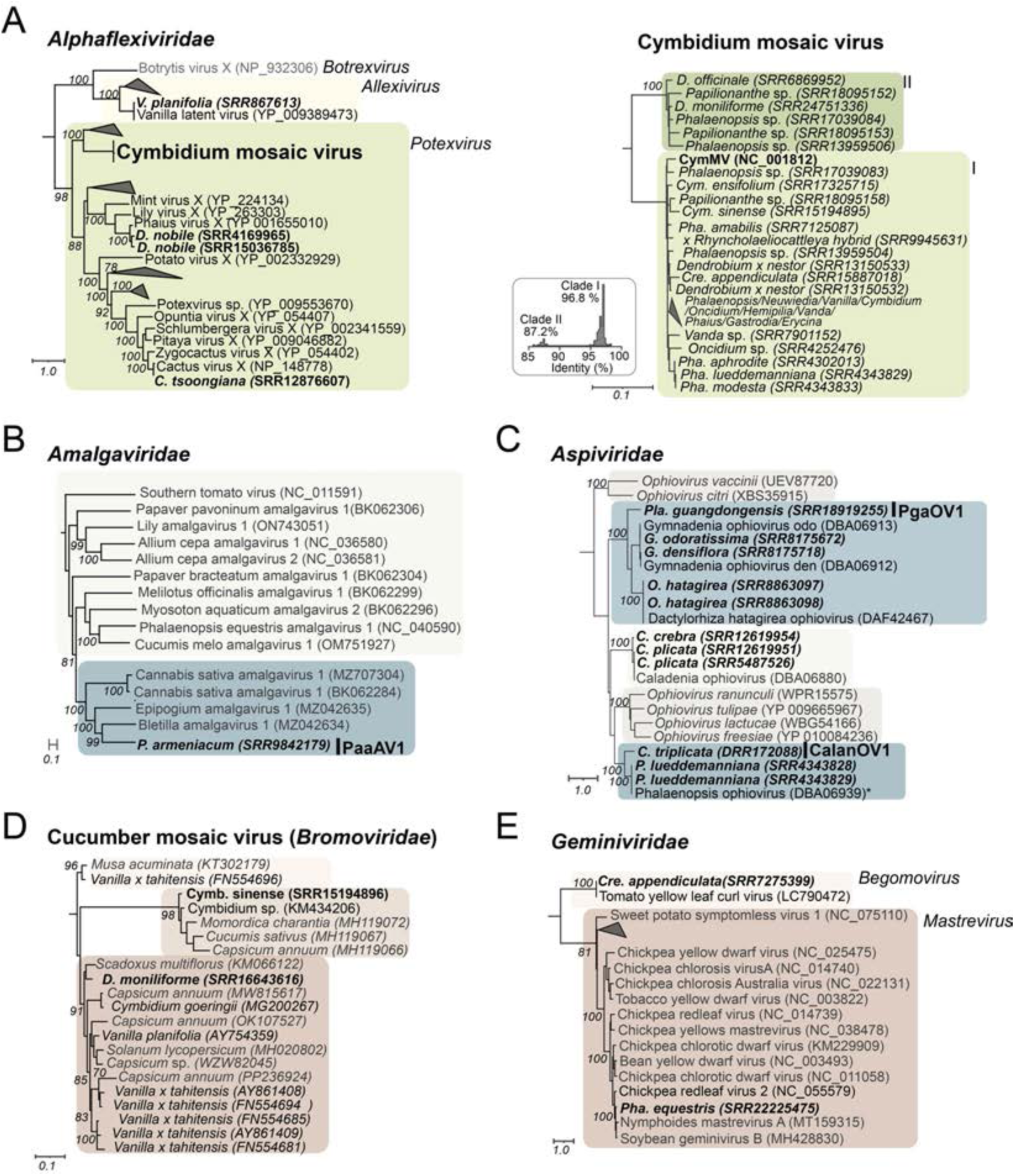
Phylogenetics trees of minor findings and known viruses associated with *Orchidaceae* RNA-seq data (Part I). Maximum-likelihood trees of representative sequences illustrating findings within *Alphaflexiviridae* (A and B), *Amalgavirida*e (C), *Aspiviridae* (D), *Bromoviridae* (E), and *Geminiviridae* (F). With the exception of the CymMV tree, sequences derived from this work are highlighted in boldface and labeled with the orchid species and the corresponding SRA accession code. Acronyms indicate putative new species. The inset in the CymMV panel represents the distribution of pairwise nucleotide sequence identities delineating clades I and II described in the text.

**Supplementary Figure 4.**
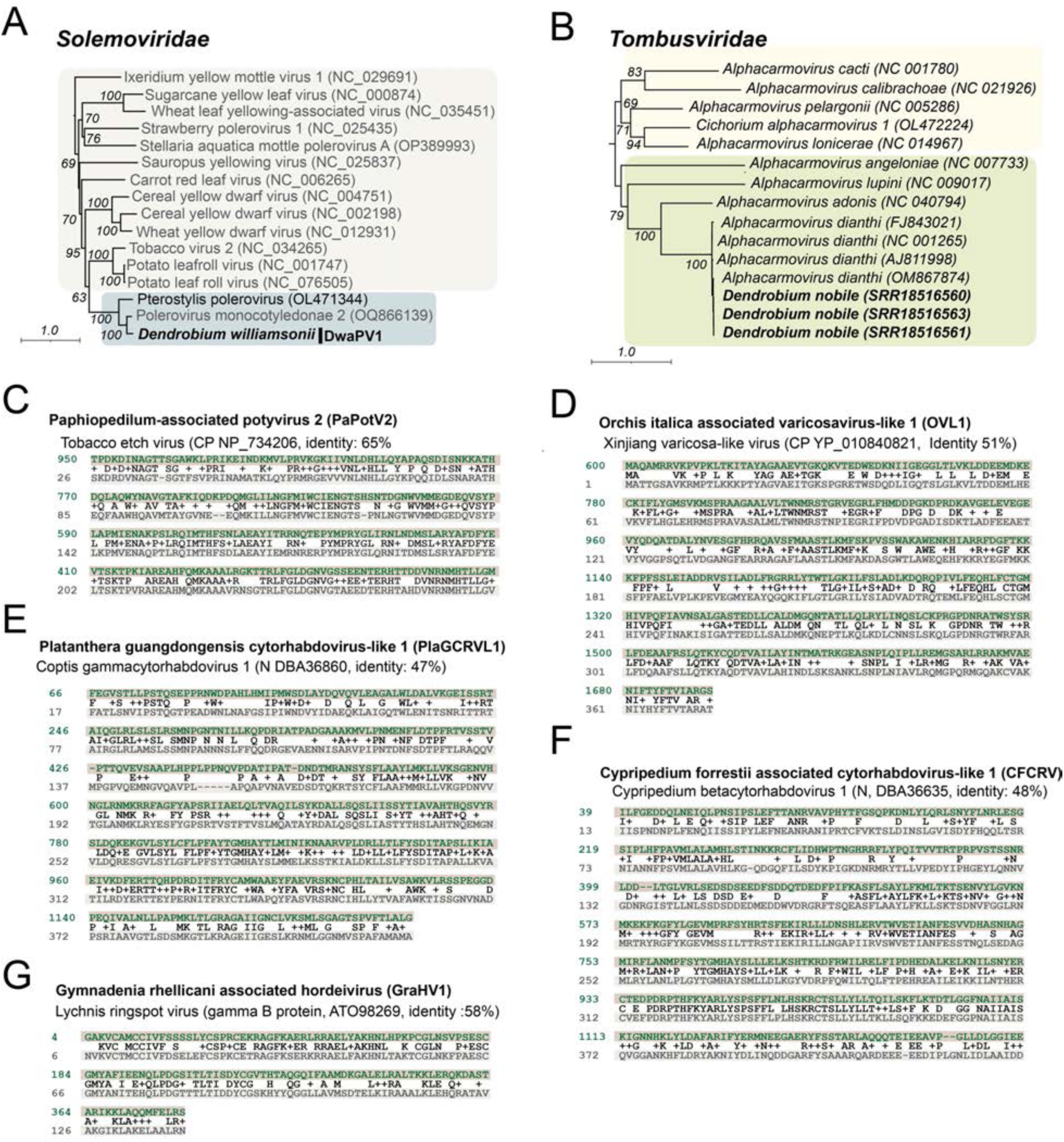
Phylogenetics trees of minor findings and known viruses associated with *Orchidaceae* RNA-seq data (Part II). Maximum-likelihood trees of representative sequences illustrate findings within *Solemoviridae* (A) and *Tombusviridae* (B). Sequences derived from this work are highlighted in boldface and labeled with the associated orchid species. Panels C to F illustrate the alignment of partial sequences that could not be included in the phylogenetic analysis but likely represent new species.

